# Optical recordings of unitary synaptic connections reveal high and random local connectivity between CA3 pyramidal cells

**DOI:** 10.1101/2025.01.15.633173

**Authors:** Robert Layous, Bálint Tamás, Arpad Mike, Eszter Sipos, Antónia Arszovszki, János Brunner, Ádám Szatai, Farah Yaseen, Tibor Andrási, János Szabadics

**Affiliations:** HUN-REN Institute of Experimental Medicine, Budapest

**Author notes:** These authors contributed equally to this work (Robert Layous, Bálint Tamás, Arpad Mike).

## Abstract

The hippocampal CA3 region is thought to play crucial roles in episodic memory functions because of the extensive recurrent connections between CA3 pyramidal cells (CA3PCs). However, different methods provided contradicting observations about the synaptic connectivity between CA3PCs. Therefore, we estimated the connectivity rate between individual CA3PCs using a new approach that is not affected by the confounds of conventional methods. Specifically, we used voltage imaging with the Voltron sensor in acute slices from rats of both sexes to test CA3PC connections by detecting spontaneous spiking and subthreshold responses in anatomically identified neurons. We detected 164 monosynaptic excitatory connections in 3078 tested CA3PC-CA3PC pairs. This 5.3% connectivity rate was much higher than that we observed with the theoretically more sensitive patch clamp method in similar experimental conditions, but it remained below the anatomically observed number of CA3PC-CA3PC contacts. We verified that the imaged excitatory connections were mediated by AMPA receptors. Our results also showed that the recurrent connections did not enrich into preferred connectivity motifs and followed a distribution that was consistent with random connectivity in general. Moreover, voltage imaging revealed CA3PCs with distinct firing properties and somatic locations corresponding to previously established heterogeneity and showed that specific connectivity rules create preferred information routes among these subpopulations. Finally, we showed that there is at least one condition, influencing patch clamp recordings but not voltage imaging, that affects the observable functional connections between CA3PCs. Altogether, our results obtained with a new voltage imaging approach argue for high local connectivity rates between CA3PCs.

**Significance Statement:** Voltage imaging offers new opportunities to measure synaptic connectivity between individual identified neurons. We employed the Voltron sensor to measure the monosynaptic connectivity between local CA3 pyramidal cells (CA3PCs) in acute slices from rats. Recurrent CA3PC connections are thought to play crucial roles in storing memory. However, the estimated connectivity rates between CA3PCs differ depending on the methods used. We showed that in spite of having a lower signal-to-noise ratio, voltage imaging results showed a much higher connectivity between CA3PCs than patch clamp recordings. Our results also suggest that the recurrent excitatory connections between CA3PCs are random, with preferred connectivity between and within subpopulations of CA3PCs.

## Introduction

One of the main features of the CA3 region of the hippocampus is the high connectivity rate between its principal cells, the CA3 pyramidal cells (CA3PCs). This extensive recurrent excitatory network was precisely described by anatomical identification of direct axo-dendritic contacts and verified synapses (Ishizuka et al., 1990; Li et al., 1994; Wittner et al., 2007; Ropireddy and Ascoli, 2011; Le Duigou et al., 2014) but it can also be inferred from the large number of dendritic spines in the radiatum and oriens regions, which are innervated only by other CA3PCs within this region. The strong recurrent excitation was confirmed in cultured slice preparations (Debanne et al., 1995). However, axonal sprouting during the culturing period could increase the number of local connections. Theoretical studies have generally agreed that abundant recurrent excitatory connections constitute an attractor network that associates random sets of neurons with specific events, and allow effective recall through partial activation (Hopfield, 1982; McNaughton and Morris, 1987; Bennett et al., 1994; Lisman, 1999; Kali and Dayan, 2000; Rolls, 2010; Li et al., 2024b). As recurrent excitation is absent in the CA1 and Dentate Gyrus regions, the CA3 is optimal for memory storage and recall within the hippocampus (Colgin et al., 2010; Kopsick et al., 2024). This consensus is further supported by the variety of cellular and molecular mechanisms, such as the different forms of synaptic plasticity among CA3PCs (Debanne et al., 1998; Lisman, 1999; Mishra et al., 2016; Li et al., 2024b). However, patch clamp studies in acute slices, which can precisely assess the functional properties of synaptic connections, have revealed unexpectedly low connectivity rates. Notably, two landmark papers reported only less than 1% connectivity rate between local CA3PCs both in rats and humans (Guzman et al., 2016; Watson et al., 2024). In contrast, a recent study in mouse acute slices demonstrated an almost 9% connectivity rate between CA3PCs (Sammons et al., 2024b). Thus, there is a contradiction between the connectivity rate observed by two gold standard methods, anatomy and patch clamp.

Recent developments in voltage imaging tools offer new approaches for studying synaptic connections. New voltage sensors can detect both the spiking activity of individual neurons and the average subthreshold synaptic responses (Abdelfattah et al., 2019; Csillag et al., 2023). We hypothesized that if patch clamp method somehow alters the apparent connectivity between CA3PCs in acute slices, voltage imaging - without the presence of patch pipettes - should reveal more local connections between individual CA3PCs in otherwise similar experimental conditions. Therefore, we imaged voltage signals in CA3 acute slices labeled with Voltron (Abdelfattah et al., 2019) to detect spontaneous spiking activity of anatomically identified CA3 neurons, and these spikes were used for measuring voltage responses in individual local neurons. We demonstrated that monosynaptic connections can be efficiently detected between identified individual neurons using optical signals alone. Even though voltage imaging is less sensitive for intracellular voltage signals compared to traditional electrophysiology, it revealed a higher connectivity rate. Furthermore, the large number of testable connections allowed us to show that the connectivity is random between CA3PCs, as predicted by theoretical studies (Rolls, 2010). Thus, our results suggest that local connectivity rates between CA3PCs are high and are better predicted by anatomical measurements.

## Results

### Patch clamp recordings and anatomy reveal different connectivity rates between CA3PCs

To replicate the contradicting observations of anatomical and electrophysiological measurements within the same slices and cells, we tested the connectivity between CA3PCs using multiple serial paired patch clamp recordings in acute slices and subsequent anatomical axonal tracing in the same acute slices. We tested single action potential (AP) or train-evoked responses in 632 CA3PC-CA3PC pairs in the CA3b and CA3c subregions. In this dataset we could identify only 3 monosynaptic excitatory connections (**Fig.1B**), corresponding to 0.47% connectivity rate, similarly to a previous patch clamp study that used similar conditions (Guzman et al., 2016). We often found disynaptic inhibitory responses between these CA3PC pairs (80 out of 581 tested, 13.8%) indicating that the axons of CA3PCs remained functional as they could efficiently excite other cell types (Miles and Wong, 1987; Beyeler et al., 2013; Sun et al., 2017). Indeed, the anatomical tracing revealed that the majority of the recorded and reconstructed CA3PCs had long axons. Intriguingly, these axons often approached and directly contacted the dendrites of other recorded CA3PCs (**Fig.1A, Extended Data Figures 1-1 and 1-2**). Between the CA3PCs, which were tested for functional connections with patch clamp pair recordings or were recorded in the same slice subsequently, we observed 93 contact sites between 626 potential pairs (14.9% connectivity rate), when we considered the strict threshold for contact sites (see Methods). Note that light microscopic analysis of biocytin labeled axons and dendrites cannot determine actual synapses, only contact sites. However, previous correlated light and electron microscopic analysis showed that the majority of putative contact sites were actual synapses (Buhl et al., 1994; Deuchars and Thomson, 1996; Buhl et al., 1997; Wittner et al., 2006). Even when every confound is considered, there is still an order of magnitude difference between the anatomical and functional prediction of CA3PC connectivity rates. Thus, we concluded that the contradiction between anatomical and electrophysiological prediction of CA3PC connectivity can be observed within the same samples and even between the same CA3PC pairs (**Fig.1C**), suggesting that the difference has a technical origin. One potential explanation is that the patch clamp pipette and recording procedure introduce physical or chemical factors that artificially alter the detectable connections between CA3PCs. Therefore, we sought to test the functional connectivity of CA3PCs with an alternative method that does not require patch pipettes at all. Specifically, we used Voltron, a new genetically encoded voltage indicator that is fast and sensitive enough to detect both spiking activity and subthreshold voltage responses from single neurons (Abdelfattah et al., 2019; Xie et al., 2021; Csillag et al., 2023; Csillag et al., 2024; Huang et al., 2024).

**Figure 1.**
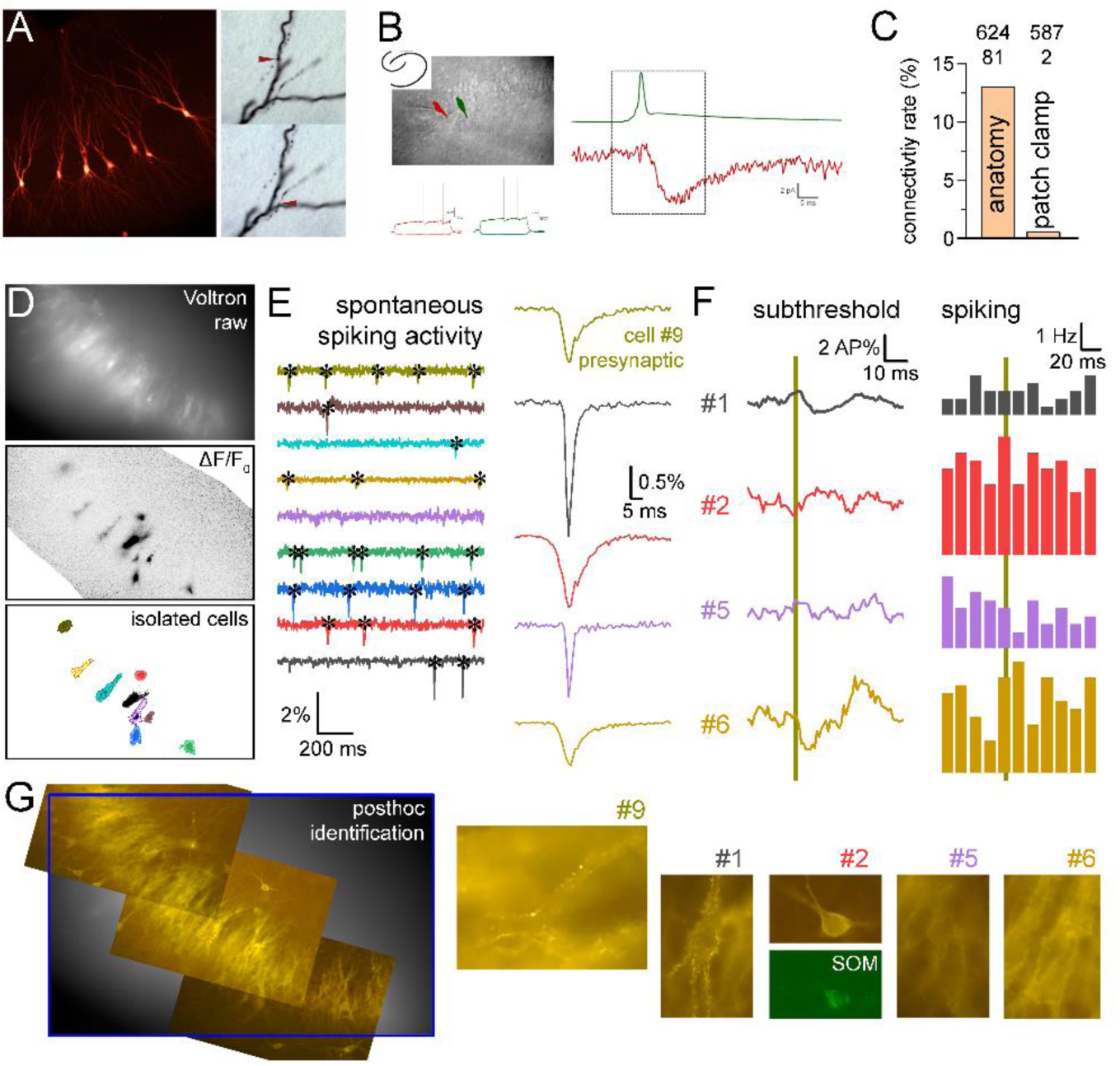
Measurements of monosynaptic connections between CA3 PCs using anatomy, patch clamp and with voltage imaging. **A.** Biocytin-labeled CA3PCs, which were tested for synaptic connections and putative contact between their axons and dendrites. More examples are shown in Extended Figures 1-1 and 1-2. **B.** Example paired patch clamp recording between two synaptically coupled CA3PCs (scale bars: 20 pA and 20 ms). **C.** Different connectivity rates revealed by patch clamp recordings and anatomical tracing within the same slices. **D.** Sparse Voltron expression (by AAVs, soma-targeted, linked with Janelia Fluor 549 HaloTag) in acute slices allowed the detection of spontaneous spiking activity in several cells, for which non-overlapping ROIs were defined. Imaging was performed at 1 kHz rate using a CMOS camera and LED light source. **E.** Individual spontaneous APs occurred randomly, and their amplitude and shape were consistent within individual cells throughout the experiments. Example traces show simultaneous signals from 9 cells and their average APs. **F.** AP timings were used to align and average subthreshold signals from other spiking neurons (whose ROI was delineated by their AP signals). These subthreshold signals (Savitzky-Golay 5-frames smoothing) were normalized to the AP peaks of individual cells (i.e. AP%) in order to obtain comparable voltage signals. Spike timings were also used for cross-correlation of activity of every cell pair to measure excitatory effects independent of subthreshold signals. **G.** Example images of the post-hoc anatomical analysis that used the preserved Voltron-JF549 staining and additional immunolabeling for somatostatin (SOM).

### Spontaneous spiking activity detected by Voltron signal can be used to test unitary synaptic responses

We injected rats (P21-23) with two rAAVs for optimal sparse expression of Voltron in CA3b and CA3c regions. The first rAAV Cre-dependently drove the expression of Voltron in the perisomatic region of neurons (AAV1-hsyn-flex-Voltron-ST), whereas the second rAAV was diluted to 1:5000 to express Cre in a smaller fraction of neurons (AAV1-hsyn-Cre)(**Fig.1D** and **Extended Data Figure 2-1**). The raw Voltron fluorescence usually did not allow distinguishing individual neurons in the dense pyramidal cell layer at the epifluorescent in situ imaging setup. However, we observed that a small fraction of neurons was spontaneously active, and their individual action potential (AP) could easily be recognized as a transient drop in fluorescence (ΔF/F_0_ images, videos or their minimal projections, **Fig.1E, Extended Data Video 2-2**). A sufficient number of active neurons (up to 31) could be identified based on the shape of their somata and proximal dendrites in ΔF/F_0_ images. We were able to identify even those neurons that elicited only a few APs during the entire experiment, or their AP gave relatively small ΔF/F_0_ signals (see Methods).

Altogether, we identified 406 spontaneously active cells in 32 slices under control conditions. CA3PCs were initially distinguished based on their wider APs, location within the cell layer, as well as thick and spiny proximal dendrites. The majority of the active cells were pyramidal cells (n = 321) consistent with their abundance relative to GABAergic cells within the hippocampus (n = 85 putative GABAergic cells) (Freund and Buzsaki, 1996). 238 of the CA3PCs and 64 of the interneurons were anatomically verified, including 31 somatostatin­expressing interneurons (**Fig.1G**). Active GABAergic cells elicited more APs during the experiments (402 ± 53 APs in interneurons vs 138 ± 9 APs in PCs); however, there was a large cell-to-cell variability in spike counts in both groups (cells with less than 10 APs were not included in the analysis). The AP amplitude of individual cells remained stable during the experiments. APs occurred apparently randomly without periodicity. Further, we did not observe a large increase in spiking activity at the onset of the imaging traces, suggesting minimal amount of photoactivated spiking was present in these conditions (Abdelfattah et al., 2019). From these observations we concluded that the consistent and reliable APs from individual neurons acquired with voltage imaging can be used as presynaptic signals for analysis of postsynaptic responses in neurons (**Fig.1F**) that were simultaneously imaged and were identified also by their spiking.

### Monosynaptic excitatory responses between CA3PCs

Not all labeled cells showed spontaneous spiking activity and, in some cases, we noticed evoked EPSP-like events in non-spiking cells. However, for the connectivity and EPSP analyses we included only spiking cells as potential postsynaptic cells for three main reasons. First, APs provided clear signals to delineate optimal ROIs for the relatively noisy subthreshold signals. Second, APs in potential postsynaptic partners allowed the normalization of subthreshold signals in each cell for its AP peak, which reduced the cell-to-cell variability of Voltron signals and provided comparable amplitude values. The third reason was that the correct calculation of connectivity rates (see below) requires that the number of potentially connected cells is exactly determined without any bias. Including only spiking cells ensured that the subthreshold Voltron signals in potentially postsynaptic cells are sufficient (i.e. good signal-to-noise) and independent (i.e. no overlap with other cells). We used strict criteria for the initial identification of monosynaptic EPSPs. First, the subthreshold EPSP-signals are expected to start at 0-2 frames (i.e. ms) relative to presynaptic AP with amplitudes of 0.2-15% of the postsynaptic AP (AP%) and to show EPSP-like waveform with faster rise and slower decay (based on raw and smoothed AP-triggered signals). If EPSP/IPSP-like events were present during baseline or several milliseconds after the presynaptic AP, the relation (both connected and non-connected) was deemed to be too noisy and excluded from further analysis. Subsequently, we used a shift window fitting algorithm with variable decay and amplitudes to verify that identified responses were above noise level (Extended Data Figure 2-1). The second criterion was that the effects of presynaptic activity on postsynaptic firing should be consistent with an excitatory effect. Note that in several cases only a few APs fell within the analyzed time window preventing a complete cross-correlational analysis. Although the signal-to-noise ratio in subthreshold signals is much lower compared to electrophysiological recordings and the number of APs for cross-correlations of spiking is low, our dual approach provided a reliable identification of excitatory connections (see below for evidence of EPSPs). However, due to the strict criteria our analysis certainly missed the smaller responses, and therefore provided only a lower bound of functional connectivity between CA3PCs.

For each EPSP-like connection we used simultaneous signals from one of the neighboring cells to verify that the event was specific to individual postsynaptic cells and not due to a fluctuation in the imaging. Control traces from neighboring cells did not show EPSP-like responses (Fig.2A). In addition to subthreshold EPSP responses, the presynaptic activity was followed by an increase in postsynaptic firing from 1.31 ± 0.12 Hz to 3.5 ± 0.37 Hz (at 4-19 ms after the peak of presynaptic AP, Fig.2B). The rise time and delay of the EPSP-like events cannot be quantified with Voltron imaging because its temporal resolution is not sufficiently fast, and because the detection of spontaneous presynaptic APs and their alignment for averaging inevitably introduce a jitter in a ±2 ms range. Nevertheless, the rise of EPSP-like events was fast, and they reached peak within 5-15 ms after the presynaptic AP. The decay time constant of the average EPSP was 8.6 ± 1.4 ms (fit on average trace) and the half-width was 17.1 ms. The average amplitude of the detected EPSP signals was 2.69 ± 0.2 AP%, with median and interquartile range 2.3 ± 2.8 % (percentage of the postsynaptic AP signal peak). Altogether, these kinetic properties are consistent with unitary, monosynaptic EPSP properties.

**Figure 2.**
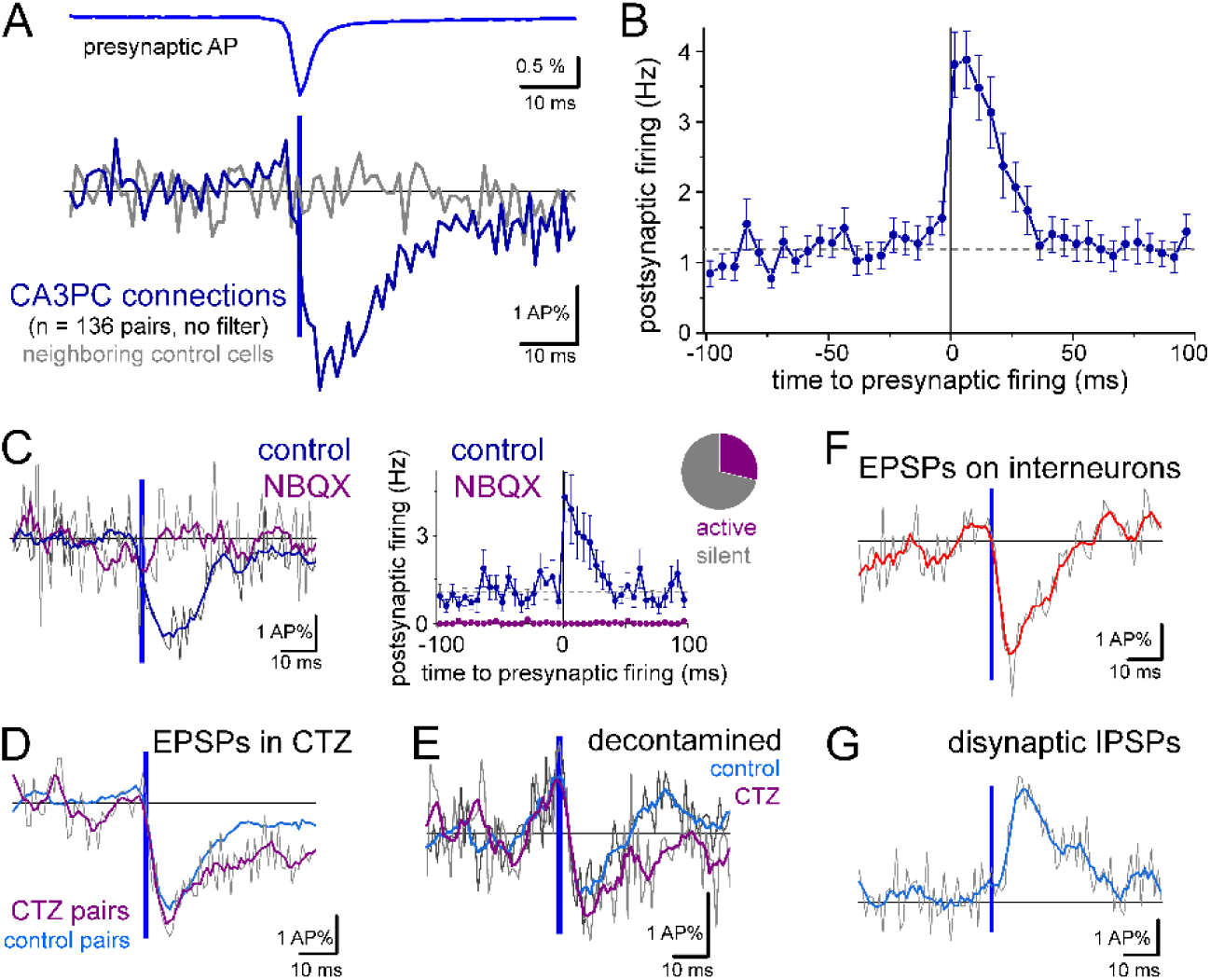
Detection of unitary excitatory postsynaptic potentials by Voltron imaging. **A.** Average of all responses identified as CA3PC-CA3PC connections (blue, n = 136 pairs from 32 slices) with average presynaptic APs (blue). As a control, gray traces show the average signals simultaneously recorded from neighboring, non-connected cells (n = 136, one for each pair). Traces were not filtered. Some pairs were excluded from this average, because of the strong contamination by postsynaptic firing. Voltron signals of each postsynaptic cell were normalized to its AP peak amplitude (AP%). **B.** Average spiking frequency of all pairs with identified connections. Only pairs with at least 20 presynaptic APs were included in suprathreshold average to avoid large relative fluctuations (n = 129 included). Zero time indicates the peak of the average presynaptic AP. **C.** Application of NBQX (10 µM) abolished responses. Slices were imaged under control conditions for the identification of connections, and then imaging continued in the presence of NBQX (n = 37 pairs). Postsynaptic traces were smoothed using a Savitzky-Golay filter (5 points, 1^st^ polynomial order) for visualization purposes (thick line, gray traces show raw signal). The majority of PCs stopped spontaneous firing after NBQX application. **D.** Average subthreshold responses and postsynaptic spiking in the presence of cyclothiazide, which inhibits AMPA receptor desensitization (CTZ, n = 68 pairs 5 slices). CTZ was present throughout these experiments. For comparison, the average smoothed response from a different set of control experiments is also shown. **E.** Average subthreshold PC-PC responses in control conditions and in CTZ, after removing traces in which postsynaptic APs occurred within the -15 ms and 50 ms time window of the presynaptic AP. This process removed only a small fraction of traces (on average 11.6 ± 1.1 traces per pair, which is equivalent 12.7 ± 0.9% of the original trace number). **F.** Average subthreshold responses in interneurons identified by their AP properties and morphology (n = 37 pairs) and in some cases (n = 11) by immunolabeling for somatostatin. **G.** Average of the occasionally observed disynaptic IPSPs between CA3PCs (n = 15 pairs).

### Pharmacological verification of EPSPs in optical pairs

To further verify EPSP identity, we employed pharmacology. The AMPA receptor blocker, NBQX (10 µM) eliminated the responses (control: 2.55 ± 0.47 AP%, NBQX: -0.33 ± 0.71 AP%, n = 37 pairs from 18 slices) and the EPSP-triggered postsynaptic spiking activities (Fig.2C). Notice that the majority of CA3PCs stopped firing after the wash-in of NBQX. Furthermore, in the presence of cyclothiazide (CTZ, 50 µM, n = 68 pairs from 5 slices, Fig.2D), which inhibits AMPA receptor desensitization, the EPSP decay was slower compared to those in separate control experiments (decay time constant: 10.5 ± 6.3 ms, half-width: 26 ms in CTZ). Although the overall firing rate was only slightly higher in the included cells, CTZ increased the number of active cells per slice that had at least 10 APs during the recorded period (18 ± 3.1 cells/slice, vs 12.5 ± 1 cells/slice in control conditions). The observed activity changes in the presence of NBQX and CTZ also suggest that spontaneous firing in the slices was mostly driven by normal synaptic connections.

We also analyzed the subthreshold responses by removing those traces where an AP occurred in the postsynaptic cells 15 ms before or 50 ms after the presynaptic APs. These de-contaminated events preserved the EPSP kinetics (both from control and CTZ pairs), but their amplitude was smaller (Fig.2E, control: from 2.7 ± 0.2 AP% to 1.5 ± 0.2 AP%, CTZ: 3.0 ± 0.3 AP% to 1.3 ± 0.2 AP%). It should be noted that these de-contaminated traces probably underestimate the average subthreshold responses, because we removed the largest events from the overall average by excluding those traces where the EPSP elicited firing.

In addition to CA3PC-CA3PC connections, we also detected EPSPs in interneurons with variable kinetics (including all types of interneurons) (Fig.2F). The decay time constant of the average EPSP in interneurons was 12.5 ± 2.7 ms and half-width: 15.5 ms. Disynaptic IPSPs between CA3PCs are prevalent under *in vitro* slice conditions and are thought to be mediated by effective excitation of intermediate interneurons (Miles and Wong, 1987; Beyeler et al., 2013; Sun et al., 2017). Thus, it requires both an intact excitatory and inhibitory connection. We observed a few clear disynaptic inhibitory responses (IPSPs) with longer delays than EPSPs (Fig.2G). However, it should be noted that the detection of chloride-mediated inhibitory events is not favored in these experiments because the resting membrane potential of cells in acute slices is close to the reversal potential of chloride. This reasoning was supported by the higher baseline firing frequency of CA3PCs with disynaptic IPSPs compared to those with monosynaptic EPSPs (2.7 ± 0.68 Hz vs 1.31 ± 0.12 Hz), which suggest more depolarized membrane potentials in the former.

Altogether, these results showed that Voltron imaging can reliably detect monosynaptic, AMPA receptor-mediated EPSPs between CA3PCs. However, the detection is likely biased towards the larger excitatory events due to the noisy signals and strict criteria used. Thus, this method probably underestimates the overall connectivity. Nevertheless, we can use this approach to estimate the lower bound of connectivity rate between CA3PCs and compare it with the theoretically more sensitive patch clamp method.

### High local connectivity rates between CA3PCs

Next, we analyzed the connectivity rates between individual CA3 neurons. For the calculation of connectivity rates, we counted not only the detected connections between CA3PCs but also counted all potentially detectable connection directions between CA3PCs. CA3PCs were identified by their spontaneous firing, soma location and shape in the fluorescent images. We excluded those potential partners where the cross-talk from the presynaptic AP signals contaminated the subthreshold fluorescent signals in adjacent cells (typically within 50 µm, depending on the orientation of dendrites). Altogether, we tested the connections between 3078 potential CA3PC pairs and identified 164 EPSP connections between them, corresponding to a 5.33 % connectivity rate (Fig. 3A). We found a similar connectivity rate when the connectivity analysis was restricted to those pairs where the identity of both the pre- and postsynaptic cells was verified as CA3PC in post-hoc anatomical identification (103 connections out of 1733 potential CA3PC connections, 5.94% connectivity rate). Furthermore, we found a similar connectivity rate between CA3PCs in independent experiments in the presence of CTZ (80 out of 1397, 5.73%).

**Figure 3.**
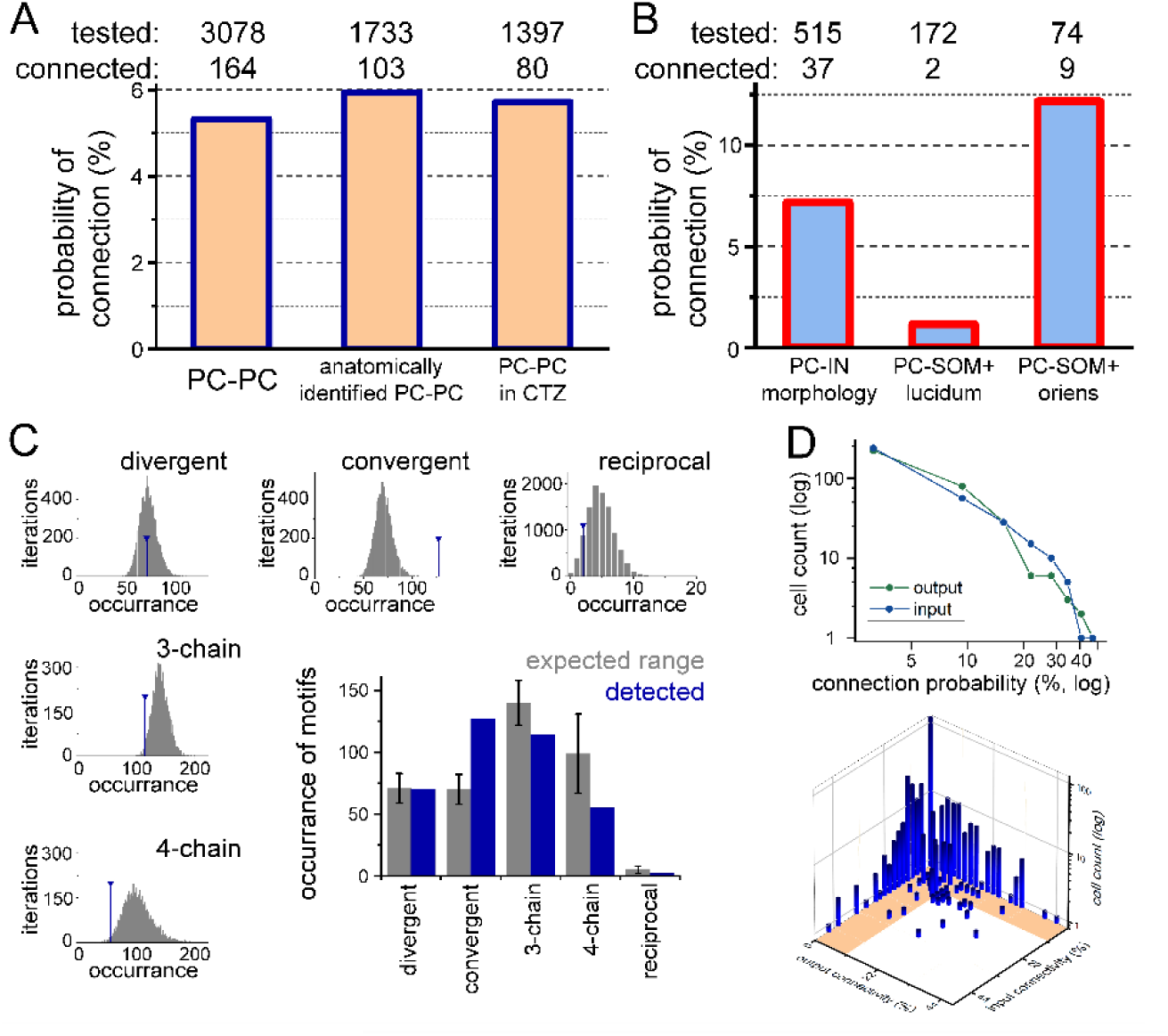
Connectivity within CA3 network. **A.** Voltron imaging revealed unitary synaptic connections between PCs. Directions were excluded both from “tested” and “connected” connections if the presynaptic AP contaminated the subthreshold signals in proximal cells, or the postsynaptic signal had unusual noise (n = 544 potential directions). The connectivity rates were not different among anatomically identified PCs and in the presence of CTZ (Fisher’s exact test p = 0.394 and 0.619). **B.** The overall connectivity rate from CA3PCs to INs. SOM+INs in the stratum lucidum were avoided by PC input (Fisher’s p = 0.011), whereas SOM+INs in the stratum oriens received input with higher probability (Fisher’s p = 0.019). **C.** Comparison of the numbers of observed and predicted connection motifs. Predictions were based on experiments with at least 9 CA3PCs, where the number of originally observed connections were assigned randomly (bootstrap iterations). Summary graph shows the predicted median and interquartile ranges (gray bars), together with the observed number of motifs (blue). **D.** The probability of forming inputs or outputs by individual CA3PCs followed a power-law distribution within our sample, but the number of CA3PCs with a large number of inputs and outputs deviated from each other. The lower graph shows the multi­dimensional distribution of input and output probabilities of individual cells. Notice that there were only 13 CA3PCs that both formed outputs and received inputs with high probability (hub cells). Shaded area on the X-Y plane indicates <11% connectivity ranges (twice the overall connectivity).

The overall connection probability from CA3PCs to anatomically identified interneurons (IN) was slightly higher than onto CA3PCs (7.18%, 37 connected out of 515 tested, Fisher’s exact test, p = 0.097). However, CA3PC-IN connection probability showed cell type specificity. Importantly, previous anatomical and physiological studies provided consistent estimations of connectivity and region-specific connectivity rates onto somatostatin­positive GABAergic interneurons (SOM+INs). We found only few connections onto SOM+INs whose soma was located in the stratum lucidum (Fig. 3B, 2 connected out of 172 tested, Fisher’s p = 0.011), which are known to be mostly hippocampo-septal projecting cells receiving excitation exclusively from dentate granule cells and avoided by CA3PCs (Wittner et al., 2006; Takacs et al., 2008; Szabadics and Soltesz, 2009). In contrast, the prevalence of EPSPs was high onto SOM+INs located in the stratum oriens (9 connected out of 74 tested, Fisher’s p = 0.019), which are mostly O-LM cells and receive extensive innervation from local PCs (Freund and Buzsaki, 1996). Thus, these results reveal a higher connectivity rate between CA3PCs than that predicted by the patch clamp approach, despite the Voltron imaging method being less sensitive. However, the connectivity rate remained below the anatomical data.

### The overall local connectivity between CA3PCs is random

The topology of connectivity matrices is an important feature of networks as they determine information routes (Barabasi and Albert, 1999; Pernice et al., 2011; Buzsaki and Mizuseki, 2014). However, similarly to the overall connectivity rates, there are conflicting predictions and observations about the structure of connectivity of CA3PCs and whether the connectivity is random (Rolls, 2010; Guzman et al., 2016; Sammons et al., 2024b). In a random network, the abundance of motifs between multiple nodes can be predicted theoretically from connectivity rates. We found that the occurrence of divergent, convergent, 3-chain, 4-chain and reciprocal motifs was different both in overall (X_2_: 18.89, p = 0.0008) and in pairwise comparisons (Fisher exact test). Therefore, we performed bootstrap analysis with 10000 iterations using all control and CTZ experiments with at least 9 CA3PCs as a skeleton, in which we randomly assigned the number of connections that we observed (Fig. 3C). This analysis showed that in cases of chain and divergent motifs, the experimentally observed number of motifs were within the predicted range. However, the experimentally observed number of reciprocal connections was lower than expected, especially compared to Guzman et al 2006, where these motifs were 6.5-times more frequent. The number of observed convergent motifs was higher in our samples than expected from random distribution. However, the higher prevalence of convergent motifs may have a technical origin because this is the only motif where two responses are measured in the same neuron. Thus, better Voltron signals in a few cells may increase the apparent number of convergent motifs with our experimental approach.

We also tested the overall connectivity probabilities at the level of individual cells, and whether the tested population follows random connectivity. In a random network - at the sample size that our measurements allowed - the numbers of occurrences (i.e. cells with certain connection probability) are expected to follow a power-law, and thus appear as a linear function on a log-log plot (Barabasi and Albert, 1999; Buzsaki and Mizuseki, 2014). We calculated input- and output-connectivity rate for each individual CA3PC in those slices where we isolated at least 9 CA3PCs (n = 322 cells in 26 slices). The results showed that the proportions of the cells with different input or output probabilities deviated only slightly from a power-law distribution (Fig.3D). Note that the total connection probabilities are expected to follow a lognormal distribution, but because of the under-sampling (tens of connections versus thousands) our measurement can detect only the upper logarithmic tail. Nevertheless, in case of non-random local connectivity, cells with high connectivity are expected to emerge from this analysis, which we did not detect, suggesting that local CA3PC connectivity is rather random, when all CA3PCs were considered (see connectivity data for distinct subtypes of CA3PCs in the next chapter).

Finally, we investigated whether “hub” cells with richer input and output connectivity exist in our sample (Morgan and Soltesz, 2008; Pernice et al., 2011). We considered a CA3PC as a hub cell if it both sent and received connections twice as likely as the overall connectivity rate (i.e. >11%). The input and output probabilities predicted 12.8 cells to fall within this category, and we identified 13 hub cells among CA3PCs. Altogether, these observations suggest that the overall local connectivity at the level of individual CA3PCs is close to random and tri-synaptic motifs are not overrepresented and reciprocal excitation is rather avoided.

### Voltron imaging reveals two types of CA3 PCs that show specific connectivity

There is a known morphological and functional diversity among CA3PCs (Hunt et al., 2018; Kveim et al., 2024) and the connectivity between these subclasses shows specificity (Sammons et al., 2024a). Recent studies categorized CA3PCs as thorny and athorny subgroups in mice based on the presence or absence of thorny excrescences, the complex dendritic spine receiving giant synapses from dentate granule cells. In most cases, Voltron labeling is not sufficient for clear identification of thorny excrescences in individual CA3PCs due to the density of labeling and perisomatic restriction of expression. Furthermore, completely athorny CA3PCs have not been observed in older rats (Raus Balind et al., 2019; Kis et al., 2024). An additional distinctive parameter of CA3PC subpopulations is the location of their soma within the cell layer as location determines availability of excitatory sources from other regions (Hunt et al., 2018) or developmental origin (Kveim et al., 2024). Athorny cells are located typically in the deep part of the cell layer (Sammons et al., 2024a). Therefore, we analyzed the connectivity of CA3PCs whose soma was in the deep, superficial or middle third of stratum pyramidale (Fig.4A). We detected more active CA3PCs in the superficial part. Similarly to observations in mouse (Kveim et al., 2024; Sammons et al., 2024a), we found that superficial and middle CA3PCs innervated deep CA3PCs with high probability, while the deep to superficial connections were the rarest (Fig.4B). However, the connectivity between CA3PCs within the same layer was not larger than the connectivity in general.

**Figure 4.**
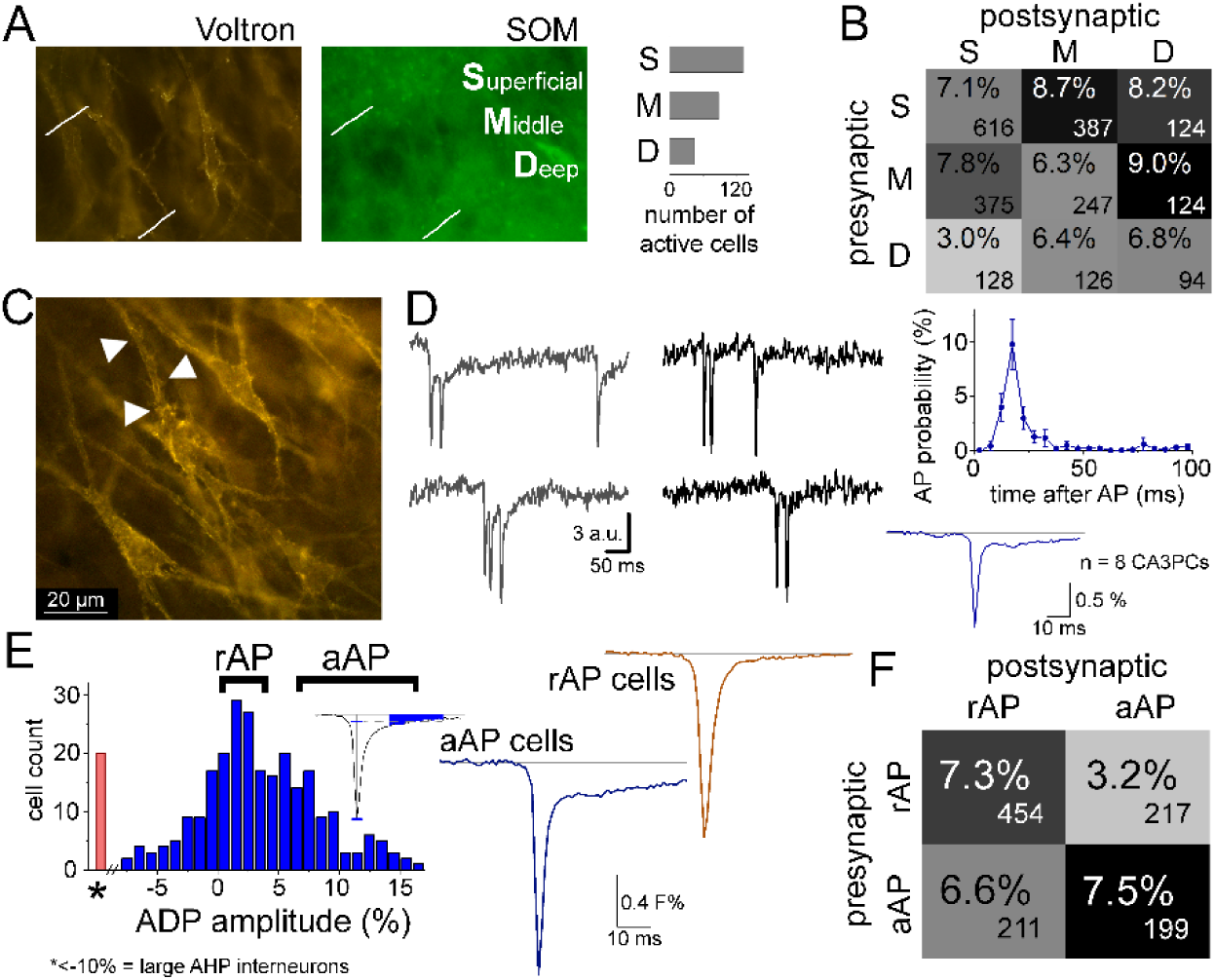
Voltron imaging reveals two types of CA3 PCs that show specific connectivity. **A.** CA3PCs were categorized as superficial, middle and deep (S, M, D, respectively) based on the location of their soma within the pyramidal cell layer. Graph shows the numbers of CA3PCs in the three categories. **B.** Probabilities and total numbers of connections between S, M, and D CA3PCs. **C.** Thorny excrescences in a post-hoc anatomical image of Voltron labeling. **D.** Bursts from two example cells (left). AP probability after first APs in 8 example cells that showed bursts (right). Average single AP shape of bursting cells. **E.** After-depolarization (ADP) was quantified in each cell (including INs with AHP) as the mean Voltron signal 15-40 ms after the AP, divided by the peak amplitude. CA3PCs were assigned into the *aAP group*, with 7-20% ADP, or into the *rAP* group, with 0-4% ADP. Traces show the average APs from 13-13 representative cells. **F.** Both aAP and rAP cells formed connections preferentially within their subgroups. Numbers of tested pairs are shown in the corners.

Based on observations from previous studies we can assume that most of the superficial cells correspond to thorny cells, whereas athorny cells were included among deep cells (Hunt et al., 2018; Sammons et al., 2024a). Although Voltron labeling was not sufficient to identify each CA3PCs as thorny and athorny, we identified 13 CA3PCs with unequivocal thorny excrescences (Fig.4C). Ten of the 13 identified thorny CA3PCs were located in the superficial layer, and 3 in the middle part, in agreement with previous observations. However, it should be noted that the identification of thorny excrescences (or their lack) in deeper cells is more difficult, because their apical dendrites pass through the usually densely labeled cell layer. Thus, we cannot unequivocally identify athorny CA3PCs. Nevertheless, deep CA3PCs were more likely to be athorny cells in mice.

CA3PC subclasses also show different firing properties. Bursting and after-depolarization (ADP) are typically observed in athorny cells (Hunt et al., 2018; Kis et al., 2024). We also observed heterogeneity at the level of AP properties, as some cells display a significant ADP, whereas the APs of other cells showed a simple exponential decay. We subdivided CA3PCs based on the ADP quantification into the *aAP* group, with a significant ADP (7-20% mean amplitude relative to peak), and into the *rAP* group, whose regular repolarization caused negligible ADP (0-4%)(Fig.4E). Similarly, to sublayer classes, aAP and rAP cells preferentially formed synapses within their groups (Fig.4F) and aAP-aAP connectivity was the highest.

Most of the CA3PCs with anatomically identified thorny excrescences had regular AP shape, and only one of them had a large after-hyperpolarization (not ADP). Bursting was rare in our sample, probably due to the low spontaneous activity rates of the CA3PCs. Nevertheless, we identified 8 bursting CA3PCs, in which at least 10% of APs were preceded by another AP within 25 ms (Fig.4D). However, we note that our analysis does not favor burst detection because of the intra-burst changes in AP amplitudes. Among the 8 bursting cells, 7 CA3PCs fell within the aAP category and showed a large ADP. These observations suggest that aAP cells are similar to athorny cells.

Importantly, our results demonstrate specific connectivity patterns within and between functional subclasses of CA3PCs. Most of our observations about the connectivity rates are in agreement with a recent study (Sammons et al., 2024a), expect for the lack of higher connectivity between deep CA3PCs.

### Effects of recording conditions on measured connectivity rates

How can the striking differences in connectivity rates from various experimental approaches be resolved? We hypothesized that the lower connectivity rate in patch clamp experiments compared to anatomy and voltage imaging could be due to technical factors related to the presence of patch pipettes filled with artificial intracellular solution. Specifically, a high concentration of ATP, GTP and K+ in typical intracellular patch clamp solutions is needed to maintain cellular health. However, during the approach of the pipette to the patched cells the intracellular solution floods the nearby tissue because positive pressure is needed to keep the tip clean. Therefore, these components that are not present extracellularly under normal conditions may activate specific pathways. For example, the activation of purinergic P2X receptors is known to selectively affect the synaptic output of CA3PCs (Wildner et al., 2024). P2X receptors are strongly and selectively expressed in CA3 axons (Sperlagh et al., 2002; Xu et al., 2016; Metzger et al., 2017; Wildner et al., 2024) and their activation opens non-selective channels that are permeable for small molecules. Other mechanisms are also activated by ATP/GTP and their hydrolytic products as they serve as a signal for tissue damage (Rodrigues et al., 2015).

To test whether the flooding of the vicinity of recorded cells with ATP and GTP influences the apparent connectivity between CA3PCs, we performed paired recordings with an intracellular solution lacking ATP and GTP, but otherwise in the same conditions as in our initial patch clamp recording. Surprisingly, we found that in these recordings the connectivity rate between CA3PCs was more than ten times higher (22 connections out of 571 tested pairs, 3.85%, compared to 0.47% in control recordings). This observation suggests that high ATP and GTP concentrations in the recording pipette have an influence on the detectability of functional connections between CA3PCs. The occurrence of disynaptic IPSPs was not changed by the removal of ATP and GTP from the patching solution conditions (56 out of 523, 10.7% compared to 13.8% control) suggesting a synapse-specific effect. The properties of the recorded EPSCs between CA3PCs matched previous data (Guzman et al., 2016; Sammons et al., 2024b; Watson et al., 2024) and were larger than the three pairs recorded in control conditions (Extended Data Figure 4-1). These results suggest that recording conditions can have substantial effects on the apparent connectivity rates between CA3PCs that may have contributed to differences between observations using various methods in various laboratories. Although further studies will be needed to elucidate the exact mechanisms underlying the specific masking of CA3PC-CA3PC connections by patching, these results reconcile the contradicting connectivity rate data with different methods.

## Discussion

Altogether, Voltron imaging experiments revealed much higher connectivity rates between CA3PCs than patch clamp recordings. However, the connectivity rate from imaging data remains below the anatomical observations, which might be due to the lower sensitivity of this new imaging method. Thus, our results suggest that functional connectivity rates between CA3PCs are close to the anatomical predictions. Anatomical studies predicted that each CA3PCs receives direct excitatory synapse from 5-20% of the local CA3PCs depending on the subregion, age and species (Ishizuka et al., 1990; Li et al., 1994; Wittner et al., 2007; Ropireddy and Ascoli, 2011; Le Duigou et al., 2014; Sun et al., 2017; Qiu et al., 2024). Based on these high connectivity rates, theoretical studies implied that CA3 operates as an attractor network, which efficiently stores memories (McNaughton and Morris, 1987; Bennett et al., 1994; Lisman, 1999; Kali and Dayan, 2000; Colgin et al., 2010; Rolls, 2010; Kopsick et al., 2024; Li et al., 2024b). Thus, our results confirm that these models with high connectivity rates provide a useful framework for understanding hippocampal functions. Unstructured or random connectivity is crucial for most theories that explain hippocampal functions at circuit level as they facilitate storing of sufficient numbers of memories and optimal recall (Kali and Dayan, 2000; Rolls, 2010). Indeed, multiple aspects of our data were consistent with random connectivity when all CA3PCs were considered. First, the connectivity of individual CA3PCs in our voltage imaging data followed power-law distribution, which is consistent with random connectivity at this sample size. Furthermore, we did not identify connectivity motifs that were more abundant than expected from random connectivity. However, our analysis suggested that reciprocal connections are underrepresented between local CA3PCs. This is in contrast to patch clamp studies which suggested either enriched reciprocal connections (Guzman et al., 2016) or similar as expected from random connectivity (Sammons et al., 2024b). The discrepancy might be of technical origin, as connection-masking effects in patch clamp may be lower in the reciprocal direction if there was already a connection, or voltage imaging may be less sensitive in the same cases. Similarly to the prediction of overall connectivity, anatomical tracing will be able to provide more direct evidence for the existence of connectivity motifs.

Our results also demonstrate that heterogeneity of CA3PCs can be observed with voltage imaging, and distinct subpopulations of CA3PCs show preferential innervation. Notably, superficial CA3PCs innervated deep CA3PCs with high probability (Kveim et al., 2024; Sammons et al., 2024a). Voltron labeling was not sufficient to identify thorny and athorny CA3PCs, a feature that is often used to delineate functionally distinct populations of CA3PCs (Hunt et al., 2018). As a note, completely athorny CA3PCs have not been observed in older rats (Raus Balind et al., 2019; Kis et al., 2024). Nevertheless, we could divide CA3PCs into functional subclasses corresponding to the thorny-athorny distinction based on other features. Namely, athorny cells were identified in the deep part of the cell layer and have been shown to elicit burst and APs with ADP (Hunt et al., 2018; Sammons et al., 2024a). In agreement with a recent study (Sammons et al., 2024a), we showed higher connectivity rates within cell groups with or without ADP, but the connectivity was less likely between different groups. Moreover, if we accept that superficial cells correspond to thorny cells while deep cells are more likely to be athorny cells (Hunt et al., 2018), our results about the innervation probability between deep and superficial CA3PCs are also consistent with the mouse data (Sammons et al., 2024a). Specifically, superficial cells innervated both other superficial cells as well as middle and deep CA3PCs with high probability. While deep CA3PCs innervated other deep CA3PCs also with high probability, they avoided superficial CA3PCs. Thus, our results support the view that there are specific connectivity rules among functional subtypes of CA3PCs. Nevertheless, further studies will be needed to elucidate further functional division of CA3PC subtypes and their interconnectivity rates because our experiments did not address subpopulations that are known to exist along the proximodistal and dorsoventral axes, or based on their different developmental origin (Sun et al., 2017; Kong et al., 2024; Kveim et al., 2024; Li et al., 2024a).

In addition to highlighting the fundamental properties of CA3 connectivity, our results also provide evidence for the utility of Voltron imaging for unitary synaptic connections in acute brain slices. Using voltage imaging has several advantages for mapping synaptic connectivity. Most importantly, it allows testing connections between a large number of neurons. Thus, a large connectivity matrix can be drawn from a single experiment. In addition to its sensitivity to single APs and subthreshold responses, the Voltron sensor is advantageous because it allows posthoc identification of neurons to which spiking or subthreshold signals from individual neurons can be assigned. We took advantage of spontaneous firing in a small fraction of neurons. Thus, we could use only optical signals to non-invasively map unitary synaptic connections. Spiking also allowed us to precisely delineate cells that we considered for analysis of subthreshold responses and morphology. The main limitation of voltage imaging compared to patch clamp recordings is the lower signal-to-noise of the signals from individual cells. Because of this limitation, our results included only the larger EPSPs between CA3PCs. Additionally, the current sensors do not favor voltage imaging of subthreshold signals with 2-photon microscopy. Therefore, this approach is limited to slice preparation before further improvements become available. Additionally, using spontaneous spiking as presynaptic signals limited us to investigate connections only between cells that fit within the field of view of our experimental setup (<1 mm). However, adaptation of our approach to stimulation of external fibers will allow extending the scope for distal connections. Even though the signal-to-noise level of Voltron imaging is not comparable to the precision of patch clamp recordings, with selective antagonist and modulator experiments we provided pharmacological evidence for the glutamatergic nature of EPSPs between CA3 neurons. Furthermore, the connectivity rates we observed between SOM+INs confirmed previous findings. An important advantage of Voltron imaging is its sensitivity and reliability. We used spontaneous spiking not only for presynaptic trigger signals, but also for identifying cells where Voltron signals are sufficiently sensitive and reliable. This is crucial for the determination of connectivity because for that not only the connected pairs should be identified but also the number of cells that can potentially be connected and these connections are available for reliable voltage measurements. Furthermore, Voltron signals from spontaneous spiking delineated the optimal ROIs for potential postsynaptic cells, allowed their post-hoc anatomical identification, and ensured that their Voltron signal was sufficient for the detection of small subthreshold responses. We also took advantage of the AP signals to normalize the subthreshold signals, as the AP amplitudes are large and similar in every neuron, allowing the elimination of variance introduced by differences in fluorescent signals (e.g. different background fluorescence). Thus, the experimental design and analysis pipeline used here will facilitate the application of voltage imaging for mapping synaptic connections. However, there are additional challenges that should be resolved for general applicability, including the restriction to in vitro or small tissue applications due to single-photon compatibility of Voltron, low number of spontaneous APs in slices, lack of absolute membrane potential values, and the limitation of the detection of chloride-mediated GABAergic events.

Our results also provide a potential explanation for the contradicting estimation of CA3PC connectivity using different methods. Specifically, our results showed that high concentration of ATP and GTP in the patching solution cell type-specifically obscures CA3PC-CA3PC connections in acute slice recordings. Thus, this effect is probably due to the high extracellular ATP or GTP concentration in the vicinity of patched cells. However, further studies will be needed to elucidate these mechanisms in detail, as the main aim of our study was to provide an estimation of CA3PC-CA3PC connectivity with a method that is not dependent on patch clamp recordings. The number of spines of CA3PCs alone predicted a high connectivity rate between CA3PCs. The only known major axon type that targets CA3PC spines in the strata radiatum and oriens originate from other CA3PCs. High connectivity rates between CA3PCs were also strongly supported by tracing studies showing electron microscopically verified synaptic contacts. Although there is a known variability depending on age, species and subregions of the CA3, the anatomically predicted connectivity rate was in the range of ten percent. Therefore, it was initially surprising that we (Fig.1B) and other research groups found very low connectivity rates with patch clamp recordings in acute slice conditions. Our observation that ATP/GTP outflow from the patching pipette potentially contributes to the loss of accessibility of the connections allows for speculation about the differences in connectivity rates observed in different laboratories. Although paired patch clamp recording is a standard method, there are several factors that change the amount of ATP/GTP in the extracellular space during the patching. For example, while larger pipettes provide better access for electrical signals due to lower resistance, they result in larger drainage of the intracellular solution. Furthermore, the number of the patched cells and pipettes in the slices can further increase the ATP/GTP levels. It is also possible that different storage methods of the patching solution contribute to different levels of excess ATP/GTP, as the half-life is within the range of typical experimental sessions at room temperature. There are other examples where a specific type of synaptic connection is particularly sensitive to slice conditions. Stable connections from cortical neurogliaform cells need extremely slow stimulation frequency and can be obtained more reliably if the synaptic activity is transiently inhibited during patching (Szabadics et al., 2007). In general, there is a good agreement in the connectivity rates revealed by the gold-standard anatomical and electrophysiological approaches in other brain areas (Buhl et al., 1994; Deuchars and Thomson, 1996; Buhl et al., 1997). We did not observe differences in the occurrence of disynaptic IPSPs, which require two synaptic steps, from CA3PC to GABAergic interneurons and from the GABAergic interneurons to CA3PCs. This suggests that these types of connections remain intact in conventional patch clamp configurations. Thus, CA3PC-CA3PC connections appear to be specifically affected by ATP and GTP in the patching solution.

## Materials and Methods

### Animals

Wistar rats (P21-23, both male and female) were used for rAAV injections in Voltron imaging experiments. The rats were bred in the Institute’s animal facility from parents originated from Charles River Laboratories breeding line. Animals were group-housed (three to four rats/cage) at constant temperature (22 ± 1°C) and humidity (40–60%), under a reverse circadian light-dark cycle. Regular laboratory chow (Sniff) and water were available ad libitum. The experiments were carried out in accordance with the European Communities Council Directive (2010/63/EU), the Institutional Ethical Codex, and the Hungarian Act of Animal Care and Experimentation (1998, XXVIII, section 243/1998) and were reviewed and approved by the National Scientific Ethical Committee on Animal Experimentation (NÉBIH PE/EA/48-2/2020). For patch clamp electrophysiology we used 23–34-day-old Wistar rats that were not injected with rAAV, but some of the patch clamp recordings and subsequent anatomy were made in slices prepared for Voltron imaging (see below).

### Voltron expression and surgery

To achieve optimal sparsity of the labeled CA3 neurons we injected a mixture of two rAAVs at four locations in each hemisphere at age of P21-23. The soma-targeted Voltron was cre-dependently expressed by one of the rAAVs (1:1.67 dilution), which was injected together with a diluted Cre-expressing virus (1:5000 dilution). pAAV-hsyn-flex-Voltron-ST was a gift from Eric Schreiter (Addgene viral prep # 119036-AAV1; http://n2t.net/addgene:119036; RRID:Addgene_119036) and pENN.AAV.hSyn.Cre.WPRE.hGH was a gift from James M. Wilson (Addgene viral prep # 105553-AAV1; http://n2t.net/addgene:105553; RRID:Addgene_105553).

The AP/ML coordinates were 4.7/±4.6 mm from Bregma, whereas the four DV coordinates were distributed between 5.3­4.4 mm. Rats were deeply anesthetized with ketamine–xylazine–pipolphen mixture (50–10–5 mg/kg) and placed in a stereotaxic frame. Viral vectors (400 nl volume/hemisphere) were microinjected at a rate of 100 nl/sec through a glass capillary (tip diameter: 20–30 μm) using a Nanoject II precision microinjector pump (Drummond). The injection pipette was left in place for 5 min to ensure diffusion before slowly withdrawn. After surgery, rats received subcutaneous injection of buprenorphine injection (Bupredine; 0.1 mg/kg) as an analgesic treatment.

### Slice preparation and dye-labeling

After decapitation under isoflurane anesthesia, 350-µm horizontal slices were cut in ice cold cutting solution (Leica VT1200S). Slices rest in a mixture of cutting and normal ACSF for 30 minutes at 33 °C. Then, we added Janelia Fluor 549 HaloTag ligand (5 nM final concentration, JF549, from Promega GA1111 or from Creative Cell, Budapest) to the solution and turned off the heating of the chamber to return the temperature to room temperature. After the 1.5 hours incubation and an intermediate washing step, the slices were transferred into a new chamber that contained room temperature normal ACSF.

### Solutions

Cutting ACSF contained: 85 mM NaCl, 75mM sucrose, 2.5 mM KCl, 25 mM glucose, 1.25 mM NaH2PO4, 4mM MgCl2, 0.5 mM CaCl2, and 24 mM NaHCO3. The normal ACSF contained: 126 mM NaCl, 2.5 mM KCl, 26 mM NaHCO3, 2 mM CaCl2, 2 mM MgCl2, 1.25 mM NaH2PO4, and 10mM glucose. All solutions were saturated with 95%/5%CO2 with constant bubbling. Imaging experiments were performed at room temperature: 23-26 °C and the slices were perfused with normal ACSF at 1-2ml/min. This lower temperature facilitated the detection of spontaneous APs in the slices. Slice holding chambers were covered with tinfoil to keep the slices in dark.

### Image acquisition

A motorized upright microscope (Eclipse FN-1, Nikon; 380FM-U, Luigs&Neumann) equipped with infrared (900 nm) Nomarski differential interference contrast optics and large field of view objective (16× NA0.8W objective, Nikon) was used to visualize slices. JF549 dye was excited with a 532nm 1W LED illumination via the epifluorescent port, 18-20mW during acquisition at the objective tip using a 519/26 – 532 - 550LP cube (Chroma). To avoid light exposure, the microscope was built within a box that covered the slice chamber (Luigs&Neumann) and the microscope up to its epifluorescent port. Light illumination and camera acquisition were triggered by an A/D converter with TTL outputs (DigiData 1440 with pClamp software, Molecular Devices). For the acquisition of Voltron signals we used a fast CMOS camera and software package (DaVinci-1K and Turbo-SM, SciMeasure Analytical Systems/Red Shirt Imaging).

First, we looked for regions within the CA3 where the labeling sparsity was optimal using low light power and short exposure. If a sufficient number of neurons was active in the selected region, we started image acquisition at 1 kHz sampling rate in a 512x320 px region (956x598 µm) in CDSBIN mode (2x2 binned, correlated double sampling). Each image sequence contained 2200 frames (2.2 seconds), which was repeated at least 100 times with 30-second resting time. Illumination started 50-200 ms before image recording. Dark frame (i.e. no light exposure) was subtracted from by the acquisition software. During the experiments the field of view and focus was corrected if there was a drift in the slice by using bright landmarks in the Voltron image. The imaging field included the complete strata pyramidale and lucidum and parts of strata oriens and radiatum.

After recording a sufficient amount of imaging data (at least 100 trials), we acquired an image video from below and above the primary z-axis at higher spatial resolution (no binning, 1024x1024 px frame, 956x956 µm). These videos helped the identification of the imaged region in the post-hoc anatomical investigations.

### Image analysis: detection and isolation of spontaneous spiking

Individual spontaneous APs could be easily identified as transiently occurring cell-shaped dark areas in the ΔF/F0 image sequences. To make sure that no APs were missed, a Python script (Py) was used to scan all image sequences, after noise reduction by an opening (erosion + dilation) algorithm. The script generated heat-maps of activity, and the active areas were re-examined in Turbo-SM to delineate initial unrefined ROIs.

For each initial ROI (provisional neuron), ΔF/F0 traces were corrected for photobleaching using a polynomial fit and exported as text files (Py). Text files were imported to pClamp, low-pass filtered at 100 Hz, and then a threshold detection function was used to find and compare APs of each provisional neuron. Thresholds were defined as at least 5 times the RMS noise of the AP-free part of the trace. Detected APs were visually verified for consistency. Cells with less than 10 APs, or with varying AP waveforms were excluded from the analysis. We observed only a few cells with abnormal transient spiking, such as sustained depolarization with high frequency initial firing and APs with decreasing amplitudes and increasing width. These cells were excluded from the analysis. The list of exact timings of APs for each individual provisional neuron was saved as a text file, which was used for further analysis. The median number of presynaptic APs was 70 (mean: 116 ± 10).

Python scripts and raw data will be available at publicly available at GitHub and ARP Research Data Repository (https://repo.researchdata.hu/dataverse/szabadics_lab).

### Image analysis: suprathreshold effects

The lists of timings were imported into an Excel template. A VBA script was used to generate cross-correlograms for all pairs of provisional neurons. For each reference cell, relative timings of APs in all other cells were plotted in the -100 ms to +100 ms window using 5 ms time bins. Cross-detection between neighboring cells was observed as peaks at zero on the histogram. In such cases the ROIs in Turbo-SM and thresholds of detection in pClamp were readjusted, and the generation of lists of timings (as described above) was repeated, until each AP was assigned to only one of the cells. Increased probability of firing within the 0 to +20 ms window indicated the possibility of synaptic connection between the reference and the target cell (if the reference cell was a PC).

### Image analysis: subthreshold responses

Based on the list of AP timings, AP-triggered average videos (-45 to +50 ms) were created for each cell from the whole field of view (Py). Minimal intensity projections were created for the 5 ms interval involving the peak of the AP. These minimal intensity projections reflect the average of all APs of a single cell throughout the whole experiment, therefore the exact, refined ROIs for individual cells could be determined. ROI definitions were converted to Fiji-compatible txt format. This set of ROIs was used to create ΔF/F0 traces for all pairs of verified active neurons (Py). For each reference cell, the average subthreshold responses of all other (potential target) cells were created and imported into an Excel template. To correct photobleaching and imaging-noise, the average of traces from silent cells (excluding those with presynaptic AP, EPSP- or IPSP-like events, or AP contamination) were subtracted from the trace of each cell in response to the same presynaptic APs. AP amplitudes of each cell were defined from its own AP-triggered video, to which postsynaptic traces were normalized. An EPSP-like deflection of the averaged ΔF/F0 trace indicated the presence of a synaptic connection. We often observed deflection in cells close to the presynaptic cell and these signals had similar kinetics as the presynaptic APs. These adjacent directions were excluded from the analysis due to potential cross-talk. We also excluded potential target cells with unstable baseline, which showed EPSP-like fluctuations.

Filtering of traces was applied only for presentation but not for analysis purposes. We used Chebyshev type 2, 8-pole low pass filter, Fstop: 0.18 or 0.24. Each potential pair was inspected by at least two experimenters and ambiguous pairs were excluded from the analysis. Those relations were excluded from analysis where the subthreshold response was contaminated by a large number of postsynaptic APs. These pairs were considered neither as connected nor not connected. However, those examples were included, if only a few postsynaptic APs contaminated the subthreshold traces that did not change the overall properties, such as kinetics.

We used a shifting window fitting algorithm for verifying EPSPs (Extended Data Figure 2-1H). An idealized EPSP waveform was repeatedly fitted to the average trace for each pair, and the RMSE of the best fit was calculated for each frame. Fits were repeated in every frame in a range of 17 frames before and 17 frames after the presynaptic APs with 1 frame-shifts (1 frame is equal to 1 ms). The parameters were constrained so that the amplitude had to be between 1-20% of the postsynaptic AP (AP%), the rise time was set to 3 ms, the time constant of decay had to be between 8-22 ms, and -1 AP% to +1 AP% was allowed for baseline shift. Mean and SD values of RMSE were calculated from the first 16 and the last 17 fits (well before or after the presynaptic AP). Two initially identified connections were dropped by this analysis, as the RMSE value did not drop after the presynaptic AP.

### Post-hoc morphological analysis after Voltron imaging

Following Voltron imaging, the slices were fixed in 0.1M phosphate buffer (pH 7.4) containing 2% paraformaldehyde and 0.1% picric acid for at least 20 hours at 4°C. The fixed samples were then resectioned to 65 µm using a Leica VT1000S vibratome. An immunofluorescence protocol was carried out for the anatomical identification of Voltron-expressing, somatostatin-positive interneurons and intrahippocampal boundaries. Voltron signal usually remained without additional staining, but it was possible to re-stain by incubating the slices in 5 nM JF549. The sections were blocked in 0.03% Triton X-100 in Tris-buffered saline with 10% normal horse serum. The sections were incubated with a rabbit anti-somatostatin-14 antibody (1:1000, BMA Biomedical, T-4103, RRID:AB_518614) for 65-72 hours and followed by subsequent incubation with dye-conjugated (AlexaFluor488) secondary Donkey antibodies. Sections were mounted in VectaShield and examined with an epifluorescent microscope. Light exposure was reduced during the anatomical processing to avoid the bleaching of the JF549 signal.

To help the identification of cells by their off-focus compartments, after the activity-imaging, a video was made that spanned in the Z-direction around the original field of view. The voltage imaged region was identified by aligning characteristic cells on the raw Voltron image and the post-hoc epifluorescent images. Cells were identified based on their shape and location on the ΔF/F0 image of their spiking. To define deep, middle and superficial portions, the stratum pyramidale was trisected horizontally. The boundaries of the cell layer were identified by the somatostatin and/or Voltron stainings. Distance between the center of the pre and postsynaptic cells were measured using the ΔF/F0 stack in Fiji.

### Patch clamp recordings

For patch clamp electrophysiology we either used slices from 23–34-day-old Wistar rats that were not injected with rAAV (n = 164 slices, n = 587 tested pairs), or slices prepared for Voltron imaging (n = 14 slices, n = 45 tested pairs). Hippocampal slices were prepared from young rats using the same protocol as for the imaging experiments. After slicing, the slices were transferred to a recording chamber of an upright microscope (Eclipse FN-1; Nikon) equipped with a 40x objective (NIR Apo N2 NA0.8W, Nikon) and infrared Nomarski DIC optics (900 nm). Recordings were performed at 33–35°C using the same ACSF solution as in the imaging experiments. Two intracellular solutions were used, each containing 1 mM NaCl, 1 mM MgCl_2_, 0.05 mM EGTA, 10 mM HEPES, 2 mM Mg-ATP, 0.4 mM Na2-GTP, 10 mM phosphocreatine, and 8 mM biocytin (pH 7.25, ∼290 mOsm). The low-chloride solution included an additional 135 mM potassium gluconate, while the high-chloride solution contained 90 mM potassium gluconate and 43.5 mM KCl. Recording pipettes with open tip resistance of 2-5 Mω were pulled from thin wall borosilicate glass capillaries (OD/ID: 1.5/1.1 mm). For experiments without ATP and GTP, Mg-ATP and Na2-GTP were omitted from the solutions without replacement by additional ions.

We used sequential paired or triplet recordings between CA3PCs and tested up to 17 potential connections per slice. The CA3PC identity was also confirmed by *post-hoc* morphological analysis of the cells. Recordings were obtained using MultiClamp 700B amplifier with DigiData1440 A/D interface. To evaluate synaptic connections, postsynaptic pyramidal cells were held in voltage-clamp mode, while the presynaptic cell was stimulated in current-clamp mode to generate either a pair of action potentials or a train of APs at frequencies up to 100 Hz. With the low-chloride intracellular solution, the postsynaptic holding potential was set to -50 mV, where GABAergic responses appeared as outward currents. For the high-chloride intracellular solution, the cell was clamped at -70 mV, with additional trials recorded at 0 mV to exclude the possibility of disynaptic GABAergic connections. In some of the experiments, we used SR95531 (2.5-5 µM, Tocris) to eliminate IPSCs. The series resistance (7-35 Mω) was monitored regularly either with small current or voltage steps in each trace. EPSC parameters were quantified by analyzing the response to the first action potential in the stimulus train, based on the average response aligned to the peak of the presynaptic action potential.

### Morphological analysis of synaptic contacts between CA3PCs

Potential contact sites between the axons of individually labeled pyramidal cells were analyzed by two staining methods. Conventional nickel-intensified DAB staining with transmitted light imaging allowed the surveying contact sites from many axons in a large area that can be traced back visually to their parent soma. Whereas, fluorescent labeling allowed high resolution imaging of potential contact sites, which were verified with immunolabeling with synaptic markers. However, tracing the axons required long additional imaging and axonal reconstructions. The two datasets were merged because we found similar connectivity rates with the two methods (41/277 and 52/349 contact sites/possible connections). Moreover, anatomical tracing data in slices from younger, non-injected animals (P23-P34) and older, Voltron-labeled animals were pooled as the overall connectivity was similar (62/403 and 31/223 contact sites/possible connections). After the patch clamp recordings, slices were fixed and re-sectioned similarly in both methods, and as in Voltron experiments described above.

*Fluorescent labeling*: Biocytin-labeled CA3PCs (2-9 cells/slices) were visualized using Alexa594 or 488-conjugated streptavidin (1:500, 1:1000, Thermofisher, S11227, S11223). To identify the possible contact sites, sections were incubated with bassoon (Bsn) antibody for 65-72 hours (mouse 1:3000, Abcam, ab82958, RRID: AB_1860018). Fluorescent dye-conjugated secondary Donkey antibodies (AlexaFluor488 or AlexaFluor594, Jackson IR, 715-545-151, 715-585-151) were used overnight. Sections were mounted and covered with Prolong Glass mounting medium (ThermoFisher). To track the local axons of multiple CA3PCs, partial reconstructions were made by Neurolucida software (version 2020.2.2, MBF Bioscience LLC, RRID: SCR_001775) using confocal image stacks (Nikon C2 scanning microscope). Initial image stacks were acquired at a resolution of 0.21 μm/px (XY directions) and 0.3 μm (Z direction) using a 60x objective (NA1.4), which was followed by higher resolution scans at potential contact sites (0.08 μm/px XY and 0.125 μm Z directions). The high-resolution images were deconvolved (Huygens software). 3D contact analysis was conducted using Nikon NIS-Elements software (version AR 4.13.04).

*DAB staining*: Endogenous peroxidase activity was blocked with 1% H2O2, then slices were incubated with ABC (avidin-biotin complex, 1:100, overnight) reagent (Vectastain ABC kit; Vector Laboratories, Burlingame, CA) in PB buffered 1% Triton X-100. The reaction was developed with DAB and NiCl2 for 3–10 minutes and stopped with H2O2 solution. Air dried sections on 3% chrome-gelatine treated slides were dehydrated (50%, 70%, 90%, 95%, absolute ethanol and HistoChoise (Sigma) than mounted with DPX mounting media (Electron Microscopy Sciences). Samples were examined with a conventional transmitted light microscope (Leica DM2500). Initial images with manually set z positions were acquired using 10x, 20x and 40x (NA1.1) objective and a camera (Nikon DS-Fi3). Partial reconstructions were made from Fiji generated stacks. The putative contact sites were imaged at high resolutions (100x, NA1.4) in a range of z-focus. Potential contact sites were identified as direct contact between traced axons and the dendritic spines of another CA3PC, or as a maximum of 2 µm distance between the axon and dendritic shaft. The latter threshold was included because it was not always possible to resolve dendritic spines. Note that more connections could be tested with anatomical tracing than with patch clamp within the same slices, because only 2 or 3 cells were recorded simultaneously, and the anatomical connections could be tested even in those cells, which were recorded subsequently. On the other hand, some of the recorded cells or slices did not have sufficient staining for anatomical measurements.

## Supporting information

Video of spiking neurons

