## Supplementary Figures for "Optical recordings of unitary synaptic connections reveal high and random local connectivity between CA3 pyramidal cells"

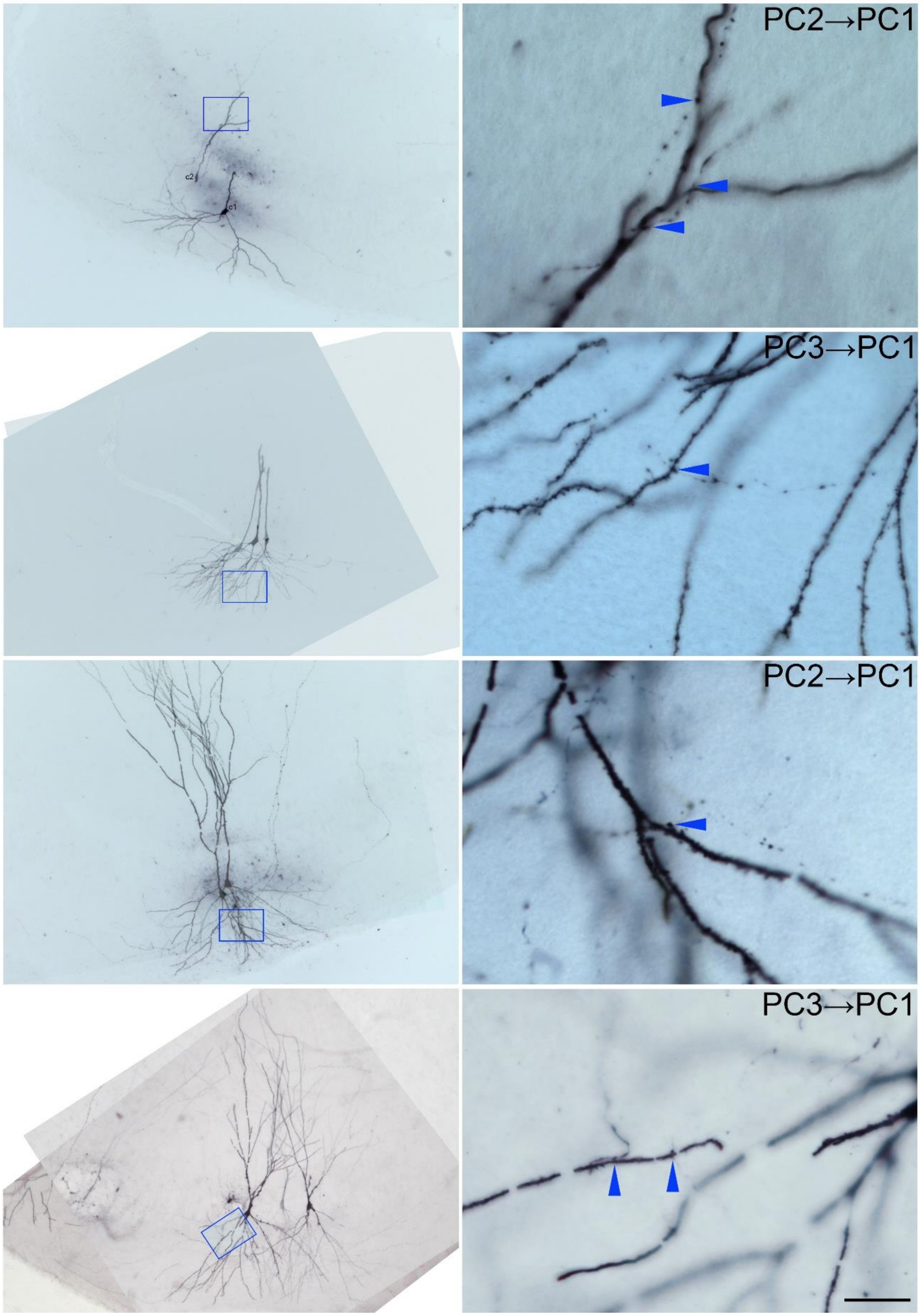


**Extended Data Figure 1-1.** Identification of contact sites between axons and dendrites of biocytin-labeled CA3PCs in DAB-stained samples. Scale bar for all high magnification images: 10 µm.

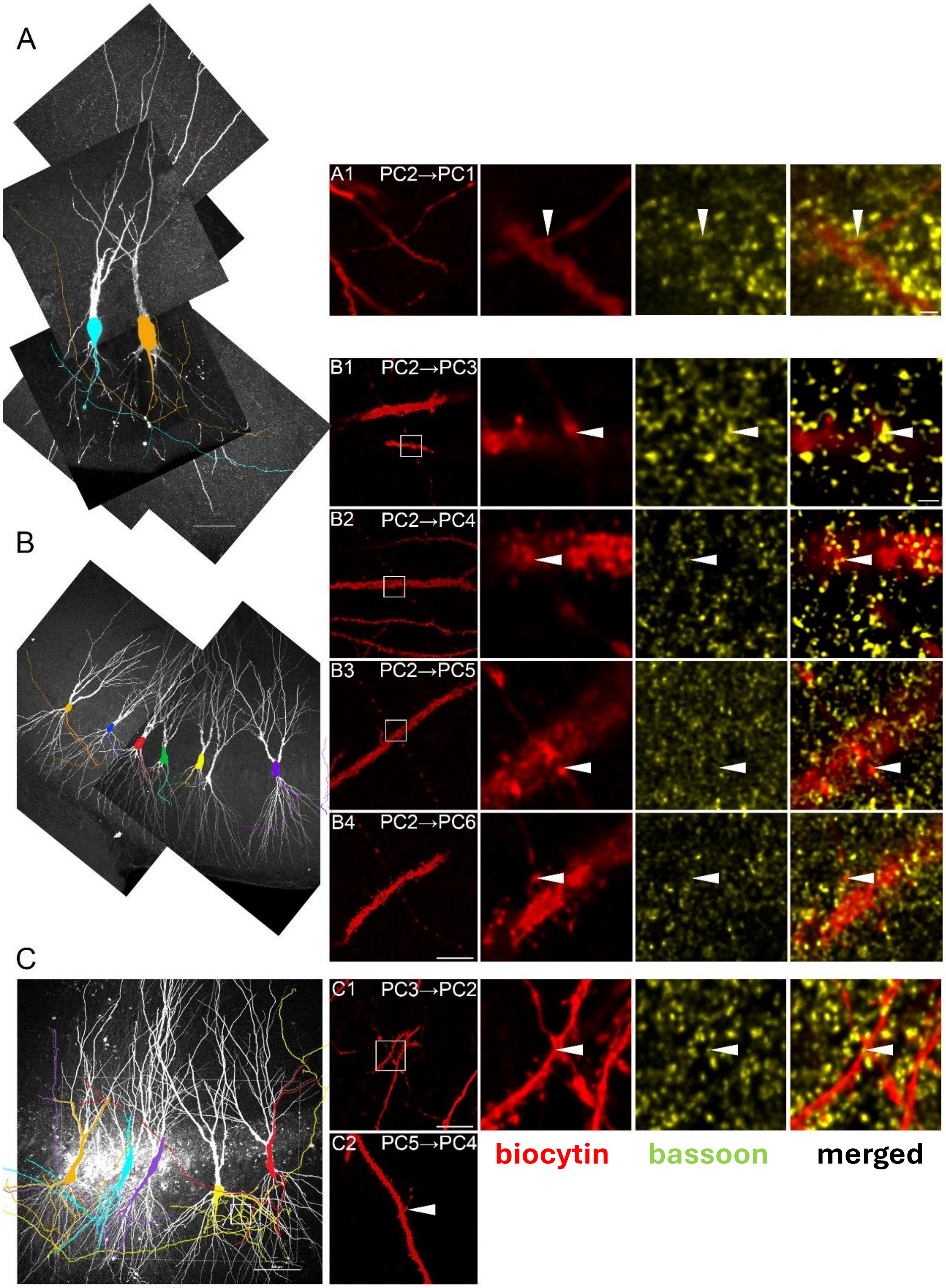


**Extended Data Figure 1-2.** Identification of contact sites between axons and dendrites of biocytin-labeled CA3PCs in fluorescence-stained samples together with immunolabeling for synaptic protein, Bassoon (green). Scale bars, for overview images: 50 μm, for maximal intensity projection images: 10 μm, for high magnification confocal images: 1 μm.

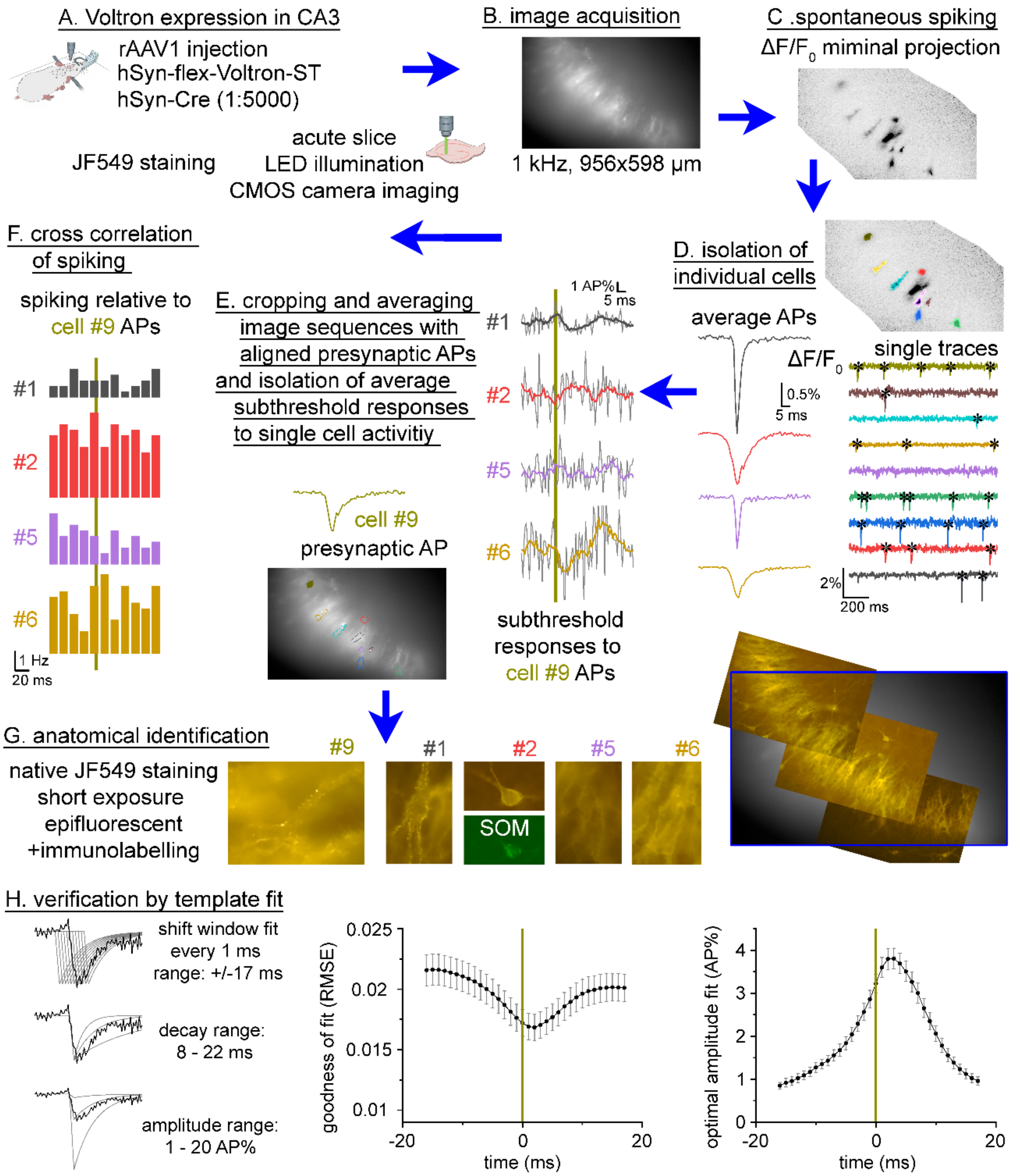


**Extended Data Figure 2-1.**

**A-G.** Workflow of detecting monosynaptic connections using spontaneous spiking and temporally aligned Voltron imaging.

**H. Shift window fitting**. Left traces show parameter ranges that were allowed to change during fitting. Each average trace was repeatedly fitted with these variable EPSP waveform in every 1 frame. Graphs in the middle and right show the average parameters obtained by the variable fits for previously identified monosynaptic connections between CA3PCs. Notice that the EPSP-waveform fit was the most optimal 2-3 frames after the presynaptic AP, as shown by the lowest RMSE and highest amplitude.

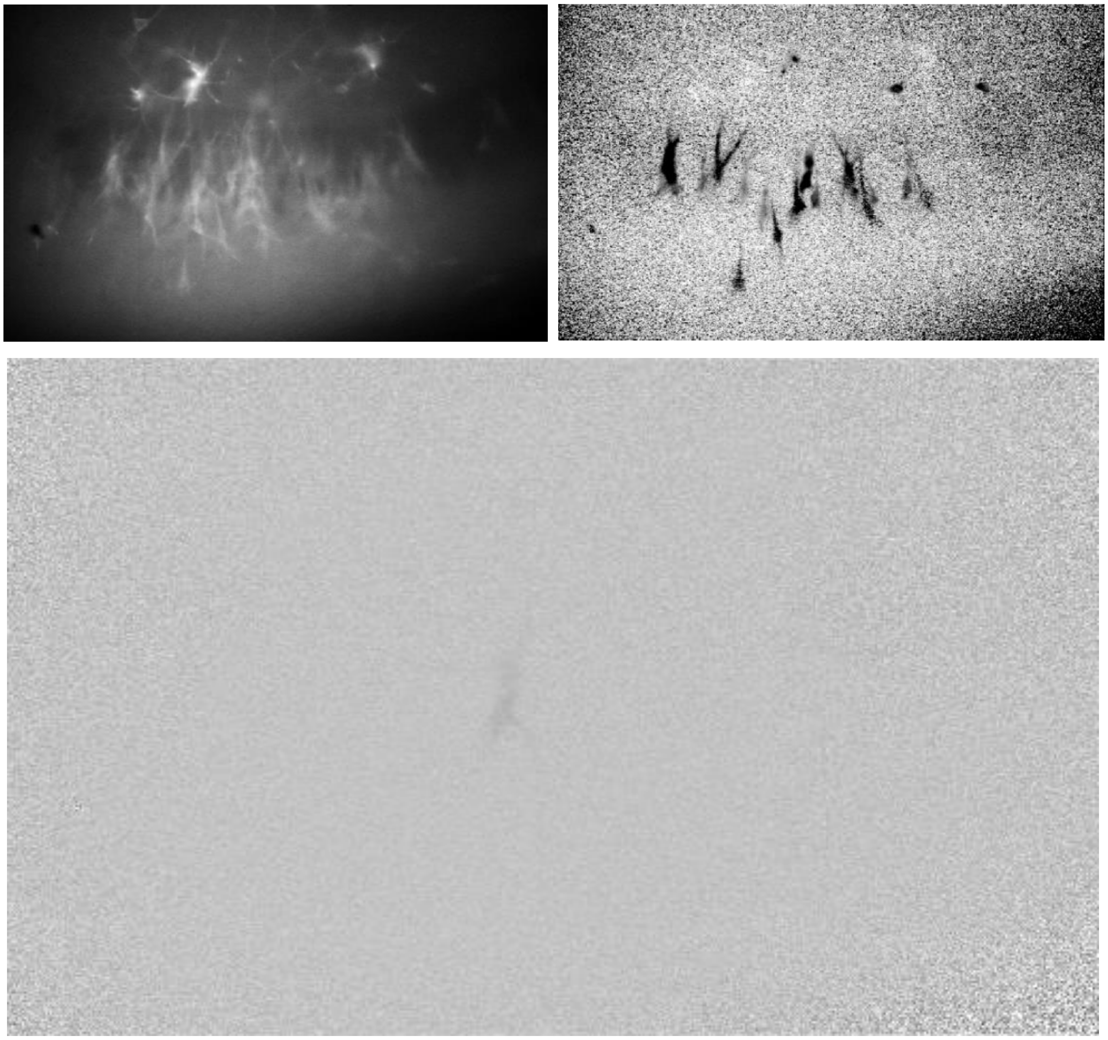

**Extended Data Video 2-2.**

**Top left:** Raw fluorescence image of Voltron-labeled CA3 neurons.
**Middle***: $\Delta F/F_0$ images of average spontaneous APs from 20 cells shown as a video.
**The PDF file shows only a single frame of the video. See separate GIF file for complete video.*
**Top right**: Minimal intensity projection of $\Delta F/F_0$ highlights all active neurons in this field view during the experiment.

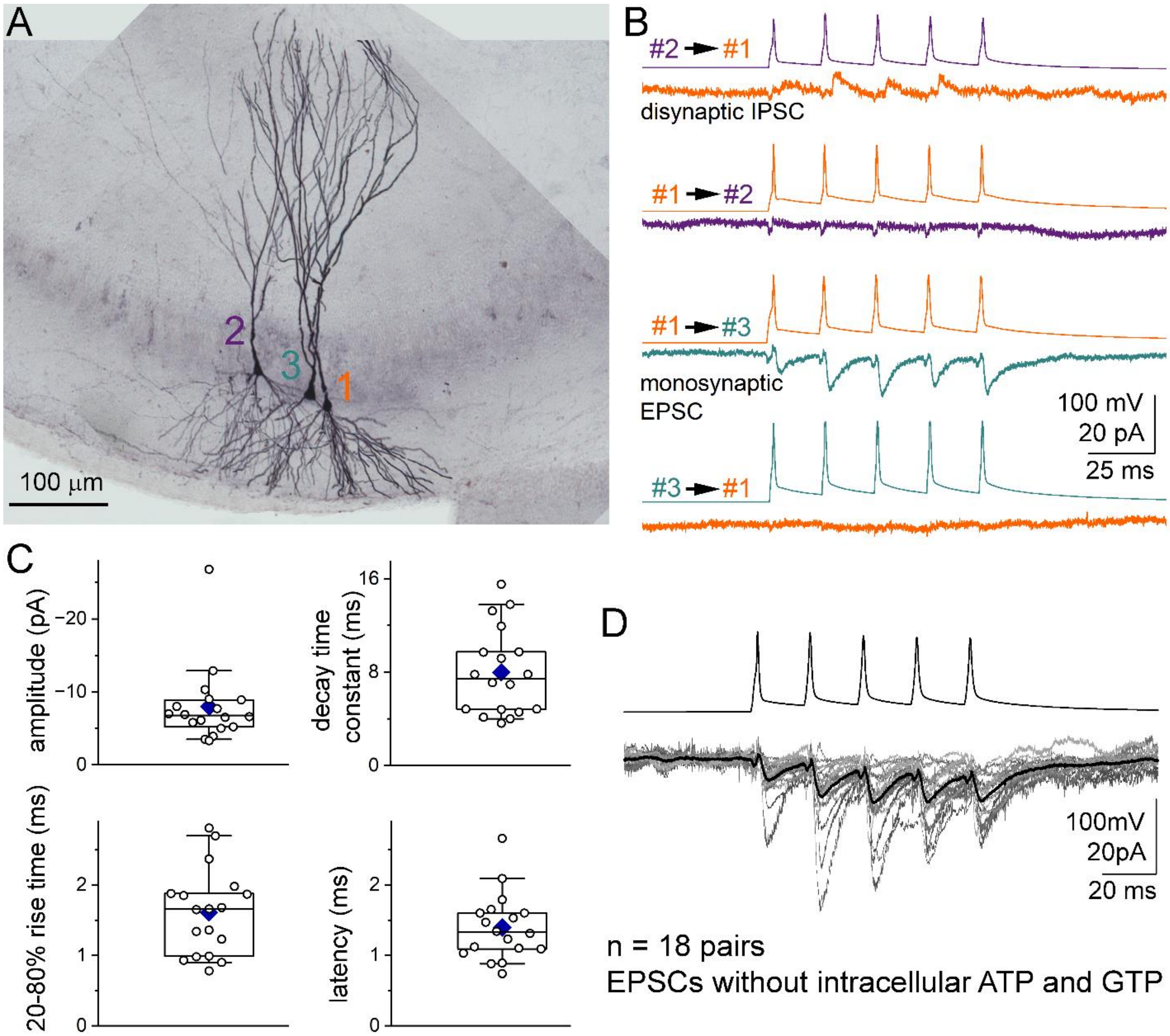


**Extended Data Figure 4-1. Electrophysiological characterization of monosynaptic connections between CA3PCs recorded without ATP and GTP.**

**A.** An example experiment demonstrating sequential paired recordings from three pyramidal cells. Collated light micrographs of biocytin-filled CA3PCs processed using DAB staining. **B.** The bottom traces show responses evoked by a short 50Hz train. Postsynaptic cells were voltage-clamped at -50 mV. In this experiment, cell #2 elicited outward disynaptic IPSC responses in cell #1 (first trace pair), whereas cell #1 evoked a monosynaptic EPSC in cell #3 (third trace pair). **C.** Summary graphs of EPSC parameters. Open circles represent data from individual connections (n = 18 pairs), blue diamonds indicate mean values, and horizontal lines denote the median value. Boxes span the interquartile range (25%–75%), while whiskers indicate the 10%–90% range. **D.** Average monosynaptic EPSCs in 18 CA3PC-CA3PC pairs recorded without ATP and GTP (gray traces) and their average (black trace). The top trace displays the average of all presynaptic AP traces.
