## Supplementary figures and images for "Optical recordings of unitary synaptic connections reveal high and random local connectivity between CA3 pyramidal cells"

### Video of spiking neurons

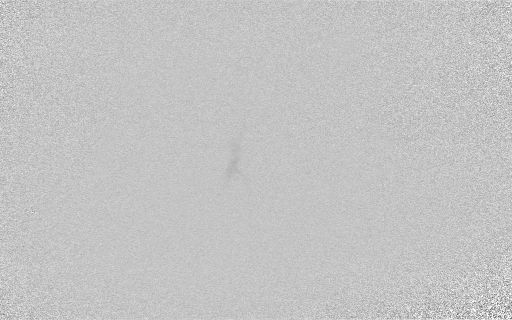
